# A signal of temporal integration in the human auditory brain: psychological insights, EEG evidence, and clinical application

**DOI:** 10.1101/2024.12.02.626219

**Authors:** Haoxuan Xu, Qianyue Huang, Peirun Song, Yuying Zhai, Xinyu Du, Hangting Ye, Xuhui Bao, Ishrat Mehmood, Hisashi Tanigawa, Wanqiu Niu, Zhiyi Tu, Pei Chen, Tingting Zhang, Lingling Zhang, Xuan Zhao, Li Zhang, Wanshun Wen, Liyu Cao, Yu Xiongjie

**Affiliations:** Department of Anesthesia, Women’s Hospital, Zhejiang University School of Medicine, Hangzhou, China; Department of Anesthesiology, Shanghai Tenth People’s Hospital, Tongji University School of Medicine, Shanghai, China; Zhejiang Provincial Key Laboratory of Precision Diagnosis and Therapy for Major Gynecological Diseases, Women’s Hospital, Zhejiang University School of Medicine, Hangzhou, China; College of Biomedical Engineering and Instrument Science, Zhejiang University, Hangzhou, China; Key Laboratory for Biomedical Engineering of Ministry of Education, Zhejiang University, Hangzhou, China; Center for Rehabilitation Medicine, Rehabilitation and Sports Medicine Research Institute of Zhejiang Province, Department of Rehabilitation Medicine, Zhejiang Provincial People’s Hospital, Affiliated People’s Hospital, Hangzhou Medical College, Hangzhou, Zhejiang, China; Department of Psychology and Behavioural Sciences, Zhejiang University, Hangzhou, China; The State Key Lab of Brain-Machine Intelligence, Zhejiang University, Hangzhou, China

**Keywords:** temporal integration, auditory cortex, click train, coma, biomarker

## Abstract

Temporal integration, the process by which the auditory system combines sound information over a curtain period to form a coherent auditory object, is essential for coherent auditory perception, yet its neural mechanisms remain underexplored. We use a “transitional click train” paradigm, which concatenates two click trains with slightly differing inter-click intervals (ICIs), to investigate temporal integration in the human cortex. Using a 64-channel electroencephalogram (EEG), we recorded responses from 42 healthy participants exposed to regular and irregular transitional click trains and conducted change detection tasks. Regular transitional click trains elicited significant change responses in the human cortex, indicative of temporal integration, whereas irregular trains did not. These neural responses were modulated by ICI length, ICI contrast, and regularity. Behavioral data mirrored EEG findings, showing enhanced detection for regular conditions compared to irregular conditions and pure tones. Furthermore, variations in change responses were associated with decision-making processes. Temporal continuity was critical, as introducing gaps between click trains diminished both behavioral and neural responses. In clinical assessments, 22 coma patients exhibited diminished or absent change responses, effectively distinguishing them from healthy individuals. Our findings identify distinct neural markers of temporal integration and highlight the potential of transitional click trains for clinical diagnostics.

## Introduction

Sound, intrinsically bound to the temporal domain, necessitates temporal integration for a coherent perception. Traditional auditory research has predominantly concentrated on the frequency domain, in which the auditory system is tonotopically organized to segregate and process distinct frequencies along the auditory pathway [1]. Nonetheless, the critical importance of the temporal dimension in auditory processing cannot be overstated. Temporal elements, including rhythm, timing, and the recognition of complex sound patterns, are fundamentally reliant on temporal cues [2]. This temporal aspect underpins essential facets of speech and music perception, alongside the discrimination of environmental sounds [3]. Temporal integration in humans spans various timescales [4], as demonstrated in oral language processing where the brain aligns with syllables, phrases, and sentences at differing temporal scales [5, 6]. Despite its significance, the mechanisms of temporal integration, particularly the direct signals related to the fusion of individual auditory units into a unified auditory object, remain largely uncharted in the existing literature.

Addressing this gap necessitates the generation of sounds with distinct components to isolate the holistic response from those elicited by individual elements. Click trains, comprised of uniform pulses that vary only in temporal spacing, serve as an exemplary stimulus for delving into temporal auditory processing. Despite their prevalent use in auditory research, click trains have rarely been studied for their holistic representation, as most research has focused on encoding individual clicks [7–9]. However, psychological studies have indicated that when the inter-click interval (ICI) of a click train is small, it can be perceived as a continuous pitch-like sound [10, 11], suggesting of temporal integration during the perception of the click train. This indicates that a click train, possessing a long-timescale structure but composed of individual clicks with short-timescale characteristics, can be perceived as a coherent auditory object rather than a collection of individual elements. However, the neural basis for pitch perception in click trains has been inadequately explored, largely due to the absence of a definitive brain signature related to pitch perception.

In our study, we employed the “transitional click train,” a stimulus created by seamlessly concatenating two click trains with differing ICIs, to explore the neural correlates of temporal integration within click trains. We hypothesized that this transitional train would trigger a change response in the human cortex, probably signifying a perceptual shift between two distinct pitches. Our results demonstrated that the human cortex indeed exhibited a specific change response closely linked to temporal integration when exposed to transitional click trains. This discovery provides a significant indicator of the holistic perception of click trains, broadens our understanding of temporal integration in auditory perception, and sets a new direction for future research in this field. Additionally, it opens new avenues for clinical applications, as temporal integration is known to be compromised in various brain disorders.

## Results

### Temporal merging into auditory objects with click trains

The human brain seamlessly integrates discrete sounds into a unified perceptual experience when the interval between these sounds is exceptionally short. For example, in the case of click trains, individuals cannot discern the gaps between clicks when the inter-click interval (ICI) falls below 29.6 ms (Supplementary Fig. 1). This phenomenon is an indication of temporal integration, whereby separate clicks merge into a single auditory object experienced psychologically. This process, where sounds with small auditory gaps integrate into a singular auditory object, is what we termed “temporal merging”. To investigate the neural representation of this temporal integration, we devised an experimental protocol featuring two types of click trains. The first type is a regular click train, characterized by uniform ICIs (the top row of Fig. 1a). The second type is an irregular click train, marked by variable ICIs (the bottom row of Fig. 1a). We created transitional trains by linking two 1-second regular click trains with different ICIs, one at 4 ms (train 1) and the other at 4.06 ms (train 2), and referred to these as Reg_4-4.06_ (the top row of Fig. 1b). Similarly, we generated transitional trains using irregular click trains with two distinct average ICIs, again one at 4 ms (train 1) and the other at 4.06 ms (train 2), and labeled these as Irreg_4-4.06_ (the bottom row of Fig. 1b). The two types of transitional trains were randomly presented to participants.

**Figure 1.**
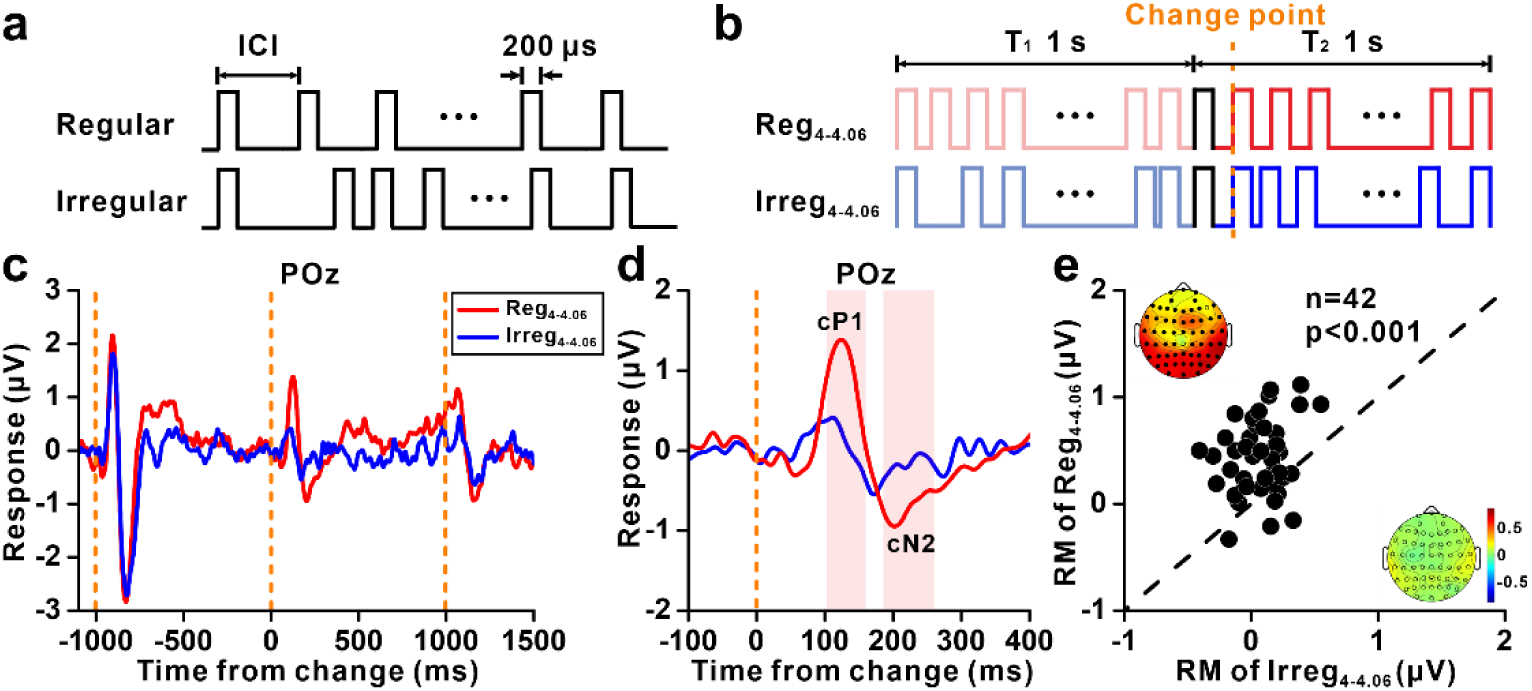
Change responses in transitional click trains. (**a**) Illustration of a regular click train (above) and an irregular click train (below), both comprising 0.2-ms pulses with specific ICI, where regular click trains maintain a constant ICI and the ICI of the irregular click train is randomized following a Gaussian distribution. (**b**) Setup of the transitional click train showing a regular transitional click train beginning with a 1-second regular click train at 4-ms ICI, transitioning into a 1-second regular click train at 4.06-ms ICI, denoted as Reg_4-4.06_, and an irregular group where regular click trains are replaced by irregular ones while keeping the average ICI equal, denoted as Irreg_4-4.06_, with the black pulse indicating the common pulse shared by both trains and the green dashed line marking the transition point. (**c**) Average responses at POz to Reg_4-4.06_ (red) and Irreg_4-4.06_ (black) across 42 subjects, with orange dashed lines indicating onset, change, and offset from left to right, respectively. (**d**) A magnified view from (c) focusing on the window from –100 to 400 ms relative to the transition point in the transitional click train. The red bars represent significant difference between the two conditions (p<0.05, two-tailed permutation test). (**e**) Scatterplots of the relative response magnitude (RM) of change responses during 74-251 ms after change for each subject under Reg_4-4.06_ and Irreg_4-4.06_, showing significantly larger change responses in the regular group compared to the irregular group (p<0.001, paired t-test). Topographic maps display the relative RM at channel level under Reg_4-4.06_ (top left) and Irreg_4-4.06_ (bottom right). Channels with significant change responses are marked with filled circles (p<0.05, paired t-test).

In the example electrode POz (Fig. 1c-d), the change responses within the time windows (92 to 148 ms and 163 to 280 ms relative to the change point) were significantly stronger for Reg4-4.06 than for Irreg4-4.06 (p<0.05, cluster-based permutation test). The change response to Reg4-4.06 in the temporal-parietal-occipital scalp regions consisted of a positive component peaking around 120 ms after the change, followed by a negative component peaking around 200 ms (Fig. 1d). To quantify this change response across the entire scalp, we designated the positive component as “change P1” (cP1) and the negative component as “change N2” (cN2), and calculated the root mean square (RMS) of the cP1-cN2 responses for each channel. Furthermore, among the 42 subjects, the majority exhibited higher response amplitudes in the time window of the change response for Reg4-4.06 compared to Irreg4-4.06 (p<0.001, paired t-test, Fig. 1e), highlighting the differential impact of regular versus irregular transitional trains on auditory processing. Across the 64 channels of the 42 subjects, pronounced change responses to Reg4-4.06 were observed in the temporal, parietal, and occipital electrodes (Supplementary Figs. 2a, c). In contrast, the response to Irreg4-4.06 primarily manifested as an onset response, with a weak or indistinct change response observed (Supplementary Figs. 2b, d).

To ascertain whether the observed change response stemmed from true temporal integration or was merely a reaction to the transient change of inter-click intervals, a systematic exploration was undertaken, involving the insertion of 1, 2, 4, 8, 16, and 32 intervals (each with an interval of 4.06 ms) into a click train with 4 ms ICI (Fig. 2a). These modified click trains were then juxtaposed against the responses elicited by both the Reg_4-4.06_ transitional trains and a continuous Reg_4_ click train (with a constant 4-ms ICI). Remarkably, the introduction of a single interval failed to produce any noticeable change responses (p=0.55, Wilcoxon signed rank test), as evidenced in Figs. 2b and 2c.

**Figure 2.**
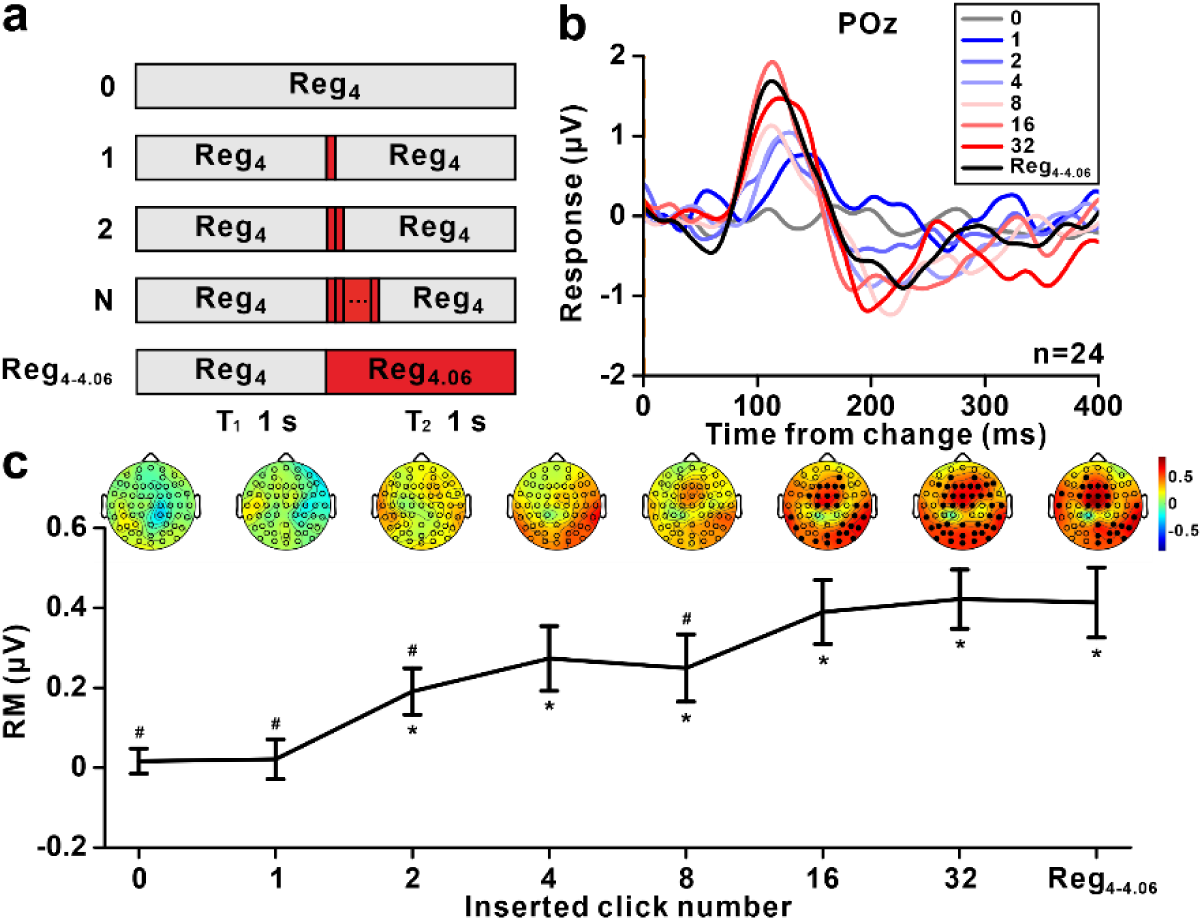
Change responses depend on temporal integration. (**a**) Experimental setup. 1, 2, 4, 8, 16, and 32 intervals, each measuring 4.06 ms, were interspersed within a click train maintaining a 4-ms ICI. These modified click trains were then juxtaposed against the responses elicited by both the Reg_4-4.06_ transitional trains and a continuous click train with a 4-ms ICI. (**b**) Average waves at POz illustrating change responses to different numbers of inserted clicks. The waves are aligned to the change point of the click train and averaged across 24 subjects. (**c**) The relative RM for change responses as a function of numbers of inserted clicks, with the time window determined with Reg_4-4.06_. The error bar indicates standard error across 24 subjects. Wilcoxon signed rank tests were made between experimental conditions and Reg_4-4.06_ (above the error bar: #, p<0.05) and between all conditions and baseline before change (below the error bar: *, p<0.05). Topographic maps (top) display the relative RM at channel level under each condition, with filled circles indicating significant change responses compared with baseline responses (p<0.05, Wilcoxon signed rank test).

It was not until the insertion of 16 intervals that the change response saturated to the level of Reg_4-4.06_ (Figs. 2b, c). Thus, the change response most likely corresponds to a perceptual shift between two pitches, coalesced through temporal merging.

### Factors affecting change response

Having established the link between the change response and temporal merging, we endeavored to delineate the requisite ICI lengths that facilitate the perceptual amalgamation of discrete auditory clicks into a singular auditory object. Employing a systematic variation of the ICI, whilst maintaining a constant ratio of ICIs between click train 1 and click train 2 (Fig. 3a), we found that an increase in ICI resulted in a diminution of the change response (Fig. 3b), with the response magnitude being inversely proportional to the ICI length (Fig. 3c). Specifically, a strong change response was elicited by both Reg_4-4.06_ and Reg_8-8.12_ (p<0.001, t-test). Notably, configurations with extended ICIs, namely Reg_32-32.48_, failed to induce discernible change responses (p>0.1, t-test), thereby suggesting that the upper limit of ICIs for the perceptual integration of individual clicks is somewhere between 16 and 32 ms, at least for the current settings.

**Figure 3.**
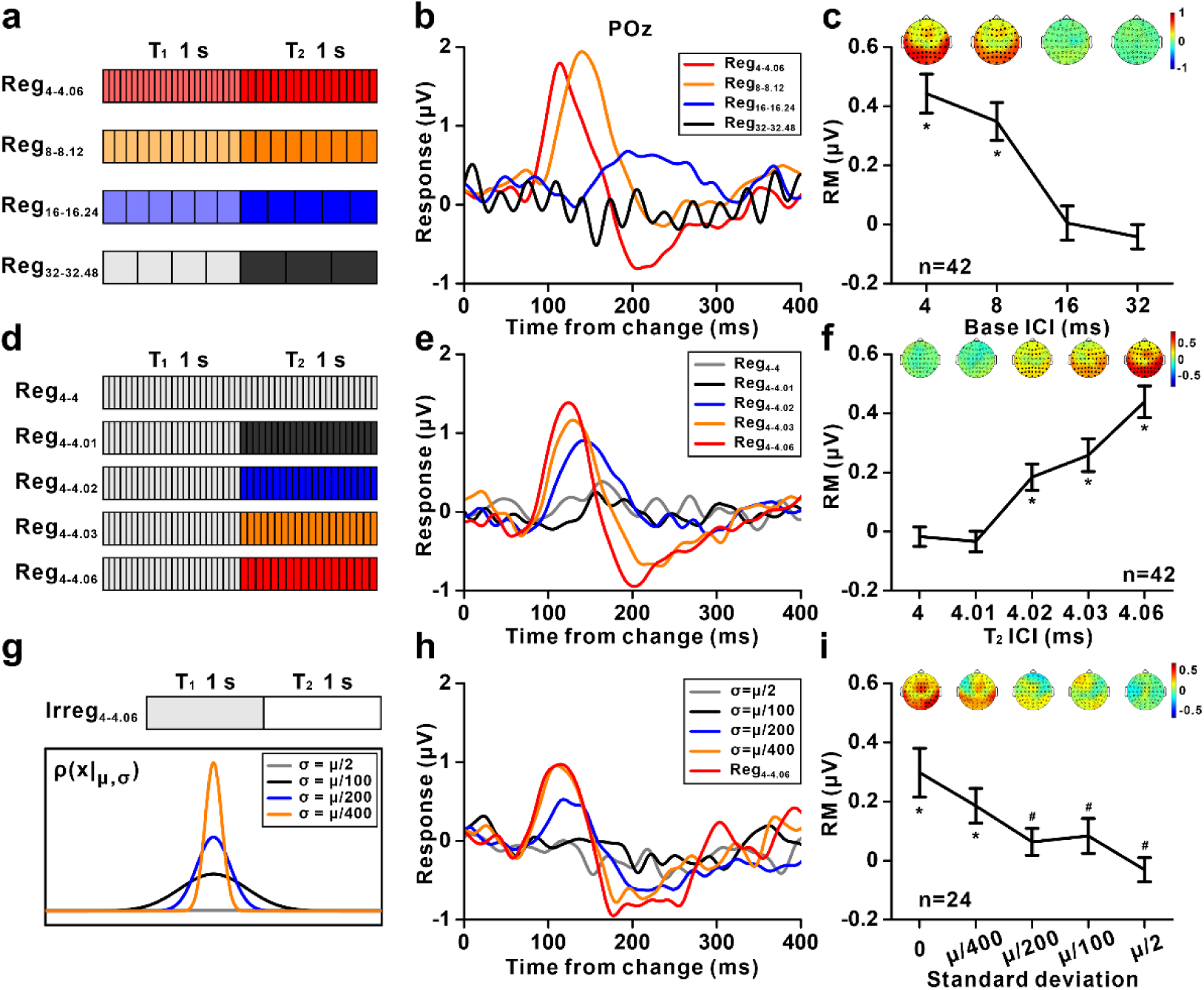
The effect of three factors on change responses in transitional train. (**a-c**) The effect of ICI length. (**a**) Stimulation setup. Four combinations of click interval were chosen in the regular transitional trains: 4-4.06; 8-8.12; 16-16.24; 32-32.48 ms. The four blocks were randomly presented. (**b**) Average change responses at POz across 24 subjects in the transitional trains for the four interval combinations. (**c**) The relative RM of change responses as a function of interval combination. Error bar: standard error across subjects. The asterisks mark significant differences (p<0.05, paired t-test) of change responses compared to the baseline before change. Topographic maps (top) display the relative RM at channel level under each condition. **(d-e)** The effect of ICI contrast. (**d**) Stimulation setup. Five levels of interval difference were chosen for the transitional trains: 4-4; 4-4.01; 4-4.02; 4-4.03; 4-4.06 ms. The five blocks were presented randomly. (**e**) Average change responses at POz in the transitional trains for the five interval differences. (**f**) The relative RM as a function of interval difference. The asterisks mark significant differences (p<0.05, t-test) of change responses compared to the baseline before change. **(g-i)** The effect of regularity. (**g**) Stimulation setup. Four levels of ICI variance were chosen for the irregular combinations in Irreg_4-4.06_: µ/400, µ/200, µ/100, µ/50 (µ=4 or 4.06 ms). Reg_4-4.06_, with ICI variance noted as 0 in (i), was also presented as a control group. The five blocks were presented randomly. (**h**) Average change responses at POz in the irregular combinations for the four levels of ICI variance. (**i**) Relative RM as a function of ICI standard deviation. Wilcoxon signed rank tests were made between experimental conditions and Reg_4-4.06_ (above the error bar: #, p<0.05) and between all conditions and baseline before change (below the error bar: *, p<0.05).

We next investigated the influence of ICI contrast on the change response. This was achieved through holding the ICI of train 1 constant (4 ms), whilst systematically modulating the ICI of train 2 so that it was longer than train 1 by a ratio between 0.25% and 1.5% (Fig. 3d). The results showed that a mere 0.5% difference in ICI (Reg_4-4.02_) was sufficient to elicit significant change responses (p<0.001, t-test, Fig. 3e), thereby indicating a remarkably high temporal resolution in the process of auditory temporal integration. A higher ICI contrast was associated with a larger change response (Figs. 3e, f).

The initial phase of our investigation predominantly focused on click trains characterized by constant ICIs, designated as regular click trains. This prompted a subsequent inquiry into the perceptual implications of varying ICIs within click trains. To quantitatively assess the effect of regularity, we introduced varying degrees of variance to each train, with standard deviations of µ/400, µ/200, µ/100, µ/2, and 0 (where µdenotes the mean ICI, set at either 4 ms or 4.06 ms, and standard deviation of 0 corresponds to Reg_4-4.06_) (Fig. 3g). Significant change responses were only observed in 0 and µ/400 conditions (p<0.05, t-test, Figs. 3h, i), with a negligible difference between the latter two (p=0.52, paired t-test). The responses with µ/400 and µ/200 were significantly weaker than those with 0 standard deviation (i.e., Reg_4-4.06_) (p<0.05, paired t-test).

### Perception performance during transitional click train

We next investigated the perception of temporal merging using behavioral experiments together with EEG recording. The primary objective was to clarify the impact of click regularity on temporal merging (Figs. 1c-e and Figs. 3g-i) and to compare the perception of temporal merging auditory objects and pure tones. For this purpose, we developed an experimental paradigm that juxtaposed three distinct sets of stimuli to assess both the behavioral performance and change responses under various degrees of contrast (Fig. 4a). The regular condition included transitional click trains transitioning from a regular click train with 4-ms ICI to another ICI (contrast levels: 0, 0.25%, 0.5%, 0.75%, 1.5%). The irregular condition comprised transitional click trains transitioning from an irregular click train (standard deviation: µ/2) with an average ICI of 4 ms to another average ICI (contrast levels: 0, 1.5%, 100%). The tone condition consisted of pure tones shifting from 250 Hz to another frequency (contrast levels: 0, 1.5%). Each block within the three conditions was designed to present a 1-second initial stimulus followed by a 1-second subsequent stimulus, concluding with a 2-second choice window. Participants were required to detect whether an auditory stimulus change had occurred (Fig. 4a).

**Figure 4.**
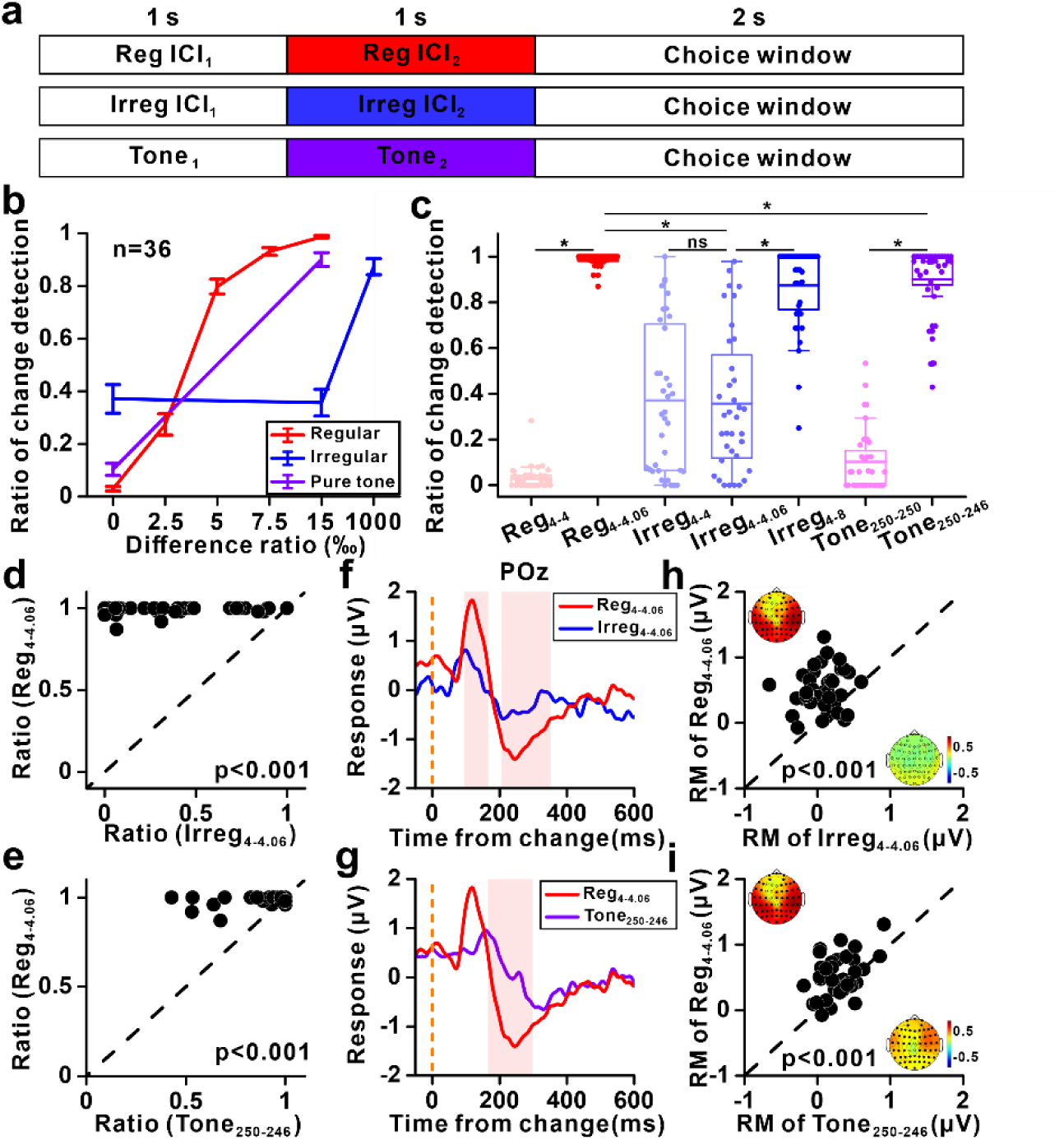
Psychological and EEG results during change detection task. (**a**) Experimental Setup outlines three sets of stimuli—regular transitional click train, irregular transitional click train, and tone changing frequency—to evaluate change response and behavioral performance under varying contrasts, featuring a sequence of stimuli presentation followed by a choice window for response. (**b**) Psychological functions depict the ratio of change detection across regular (red), irregular (blue), and tone (purple) stimuli, with standard error bars for 36 subjects. (**c**) Boxplot illustrates the change detection ratio among groups (paired t-test, with ‘*’ indicating p<0.01 and ‘ns’ indicating non-significance). (**d-e**) Scatterplots compare change detection ratios between Reg_4-4.06_ and Irreg_4-4.06_, and Reg_4-4.06_ and Tone_250-246_, respectively. (**f-g**) Average waves at POz aligned to the change point show comparisons between Reg_4-4.06_ and Irreg_4-4.06_, and Reg_4-4.06_ and Tone_250-246_, with red bars indicating significantly different time window (two-tailed permutation test). (**h-i**) Scatterplots of the relative RM of change responses comparing: Reg_4-4.06_ vs. Irreg_4-4.06_ (h), and Reg_4-4.06_ vs. Tone_250-246_ (i), where p<0.001 for paired t-tests. Topographic maps display the relative RM at channel level under Reg_4-4.06_ (h and i, top left), Irreg_4-4.06_ (h, bottom right), and Tone_250-246_ (i, bottom right), with filled circles indicating channels with significant change responses (p<0.05, paired t-test).

The results showed that the change detection performance progressed with the increase in the difference between the first and the second stimulus (Fig. 4b). Remarkably, a 1.5% contrast difference in the regular condition (Reg_4-4.06_) led to a detection rate of 98.8% in correctly identifying changes (Fig. 4c), in stark contrast to the detection rate of 35.7% observed in the irregular condition (Irreg_4-4.06_), which did not significantly differ from its control, Irreg_4-4_ (p=0.36, paired t-test, Fig. 4c). The detection rate in the irregular condition reached 87.4% when the contrast was increased by 100% (i.e. Irreg_4-8_, Fig. 4b). For pure tones, the detection rate was 90.1% when the tone shifted from 250 Hz to 246 Hz (Tone_250-246_). Note that 250 Hz corresponds to 4 ms, and 246 Hz corresponds to 4.06 ms. The detection rate for Tone_250-246_ was significantly higher than the control condition (Tone_250-250_), in which the tone was always 250 Hz (p<0.001, paired t-test, Fig. 4c), yet lower than that observed for Reg_4-4.06_ (p<0.001, paired t-test, Fig. 4c). Furthermore, subject-by-subject comparisons revealed most subjects had higher detection rate for Reg_4-4.06_ than for Irreg_4-4.06_ (p<0.001, paired t-test, Fig. 4d) and Tone_250-246_ (Fig. 4e). In summary, these findings emphasize the enhanced performance in the regular condition in identifying contrast changes compared to both the irregular condition and the pure tone condition.

For the EEG change responses in the three conditions, Reg_4-4.06_ evoked stronger change responses compared to Irreg_4-4.06_ (Fig. 4f) and Tone_250-246_ (Fig. 4g). Actually, no significant change response was observed in Irreg_4-4.06_ (Fig. 4f). Individual results also indicated that most subjects demonstrate stronger changes responses for Reg_4-4.06_ than for Irreg_4-4.06_ (p<0.001, paired t-test, Fig. 4h) and Tone_250-246_ (p<0.001, paired t-test, Fig. 4i), which is consistent with the behavior results (Fig. 4d and Fig. 4e). Interestingly, the variation in change response amplitude was correlated with decision-making in the more difficult condition, such as Reg4-4.01. In Reg4-4.01, the decision to detect a change was typically accompanied by a stronger change response compared to the decision of no change in the sound (Supplementary Fig. 3).

### The effect of temporal continuity

To investigate the impact of temporal continuity on the change responses, we designed two sets of stimuli: one set without gaps of silence between click train 1 and click train 2 (No-gap) and the other set with a gap of 600-ms silence between the two click trains (Gap). Four transitional click trains were used: Reg_4-4.01_, Reg_4-4.02_, Reg_4-4.03_, and Reg_4-4.06_. Participants were asked to detect whether an auditory stimulus change had occurred (Fig. 5a). The behavioral performance was better for the No-gap click trains than for the Gap click trains (Fig. 5b), with most participants showing this pattern (p<0.001, paired t-test, Fig. 5c).

Subsequently, we examined the EEG responses, superimposing all contrast conditions for both the No-gap (Fig. 5d) and Gap (Fig. 5e) stimuli. The responses to varying contrast conditions were distinguishable in the No-gap condition (Fig. 5d) but nearly indistinguishable in the Gap condition (Fig. 5e). We plotted the tuning curve across two windows: peak response ([70 120] ms, Fig. 5f) and trough response ([133 183] ms, Fig. 5g). As the contrast increased, the response magnitude also increased, displaying clear tuning in the No-gap condition for both windows (p<0.001 for both windows, ANOVA test, red line in Figs. 5f-g), while remaining nearly constant in the Gap condition for both windows (p>0.1 for both windows, ANOVA test, blue line in Figs. 6f-g). These EEG findings align with the psychological results regarding thresholds (Fig. 6c), emphasizing the role of temporal continuity in both psychological perception and neural processing.

**Figure 5.**
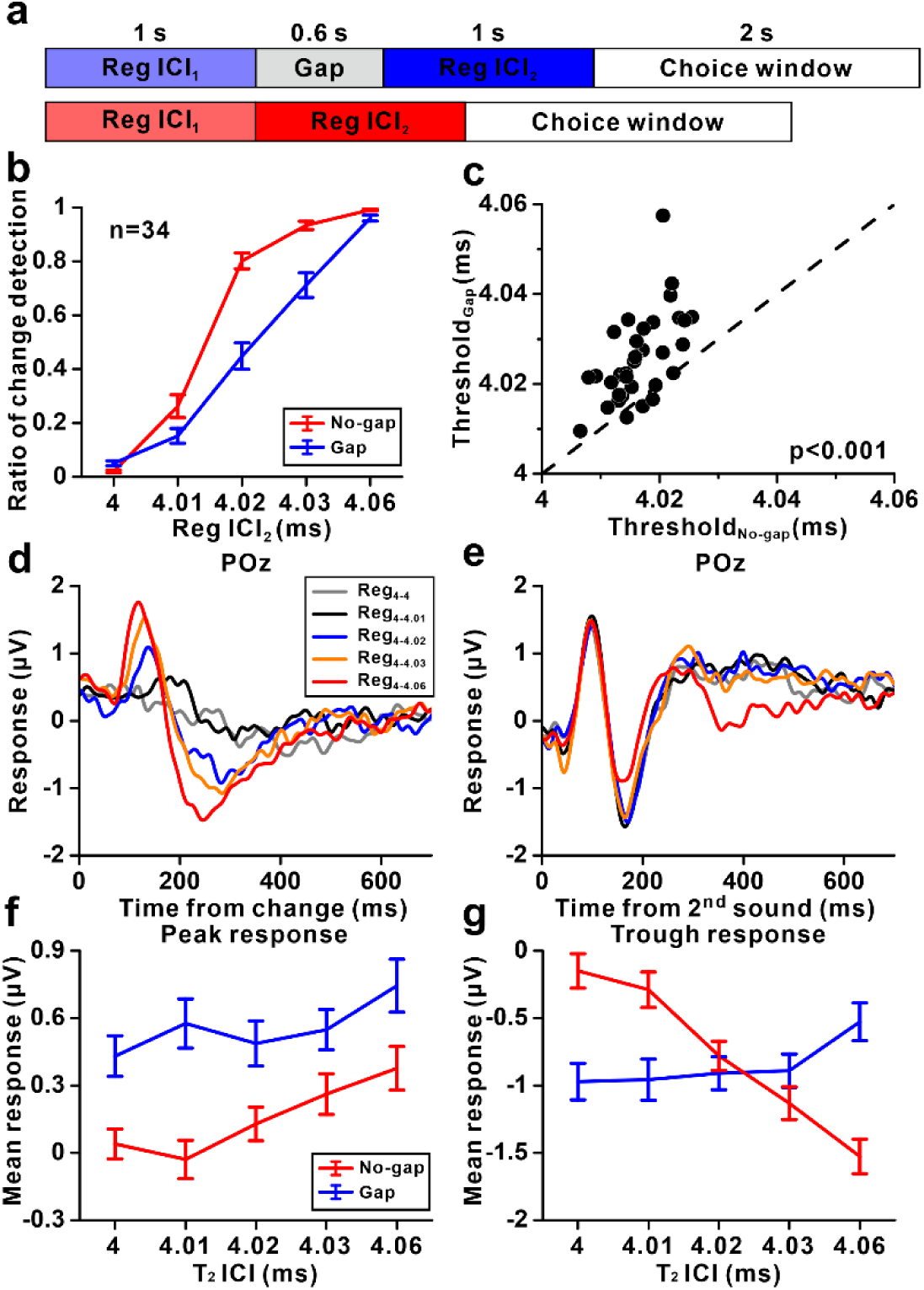
The effect of temporal continuity on psychological and EEG responses during change detection task. (**a**) Experimental setup with two stimulus sets: the transitional train (no-gap) with varying contrasts (Reg_4-4_, Reg_4-4.01_, Reg_4-4.02_, Reg_4-4.03_, Reg_4-4.06_), and a similar set that includes a 600-ms gap between click trains. Participants identified auditory changes by pressing designated buttons. (**b**) Psychological functions depict the ratio of change detection across no-gap (red) and gap (purple) change detection task, with standard error bars for 34 subjects. (**c**) Scatterplots of behavioral thresholds in the two behavioral conditions (p<0.001, paired t-test). (**d**) Average waves at POz aligned to the change point under different stimuli in no-gap behavioral section. (**e**) Average waves at POz aligned to the onset of the second sound under different stimuli in gap behavioral section. (**f**) The average peak response as a function of ICI contrast for change responses under no-gap (red) and gap (blue) conditions. (**g**) The average trough response as a function of ICI contrast under gap condition.

**Figure 6.**
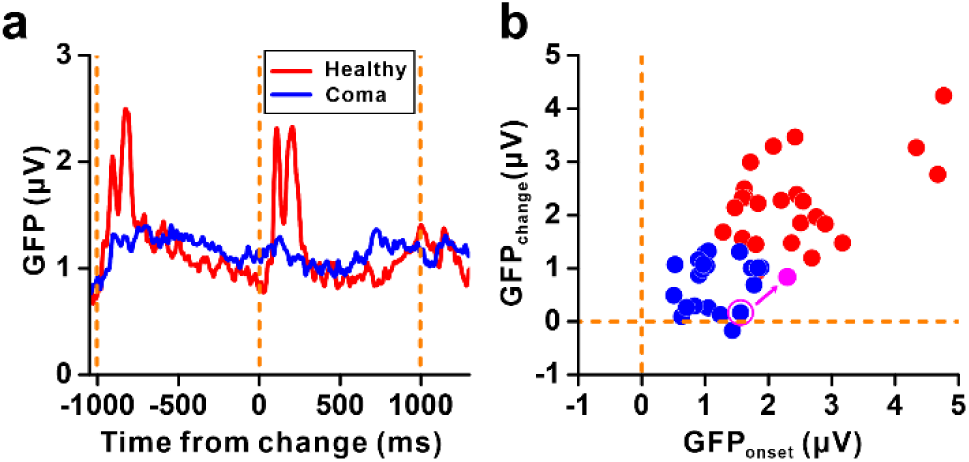
The effect of consciousness on change responses in transitional train. (**a**) Averaged global field power (GFP) of coma participants (blue line, n=22) and healthy participants (red line, n=24) under Reg_4-5_. (**b**) Scatterplots of GFP indices of change responses and onset responses for coma participants (blue) and healthy participants (red). The magenta circles indicate the subject before (hollow) and after (filled) recovery.

### Potential clinical application

Considering the fundamental role of temporal integration in the brain [12, 13] and its relevance to many psychiatric diseases [14–18], the change response serves as a promising tool for diagnosis. To explore the potential for clinical application of this paradigm, we conducted 64-channel EEG recordings in 22 coma subjects using transitional click trains stimuli: Reg_4-4_ and Reg_4-5_.

For coma patients, both onset and change responses were small, even in Reg_4-5_ (Supplementary Fig. 4a), contrasting with the healthy subjects (Supplementary Fig. 4b). The scatter plots of onset vs. change responses showed significant overlap between coma subjects and healthy subjects. This overlap was probably due to the presence of slow oscillations with larger amplitudes localized to specific channels, along with the prolonged latency and extended duration of the auditory response in coma subjects (Supplementary Figs. 5a-c). To quantify change responses in subjects with impaired consciousness, global field power (GFP) was employed due to its robustness to spatial variability and enhanced sensitivity to response latency [19, 20]. GFP calculates the standard deviation of EEG data across all electrodes at each sampling point, thereby mitigating the influence of spatial variability in electrode placement or individual differences in brain anatomy. This is crucial when studying subjects with impaired consciousness, where localized differences in brain function might occur due to injury or pathology. A robust onset response was detected in one example coma subject using GFP (Supplementary Fig. 5c), although no visible onset response was detected in amplitude (Supplementary Fig. 5b). However, no change response was detected even using GFP (Supplementary Fig. 5c). In the population, no visible onset or change responses were detected using GFP in coma patients, whereas significant robust responses were observed in healthy subjects (Fig. 6a).

Furthermore, the scatter plot of onset vs. change responses in GFP effectively separated coma patients from healthy subjects, suggesting the transitional click train paradigm as a good tool for distinguishing between the two groups (Fig. 6b). More interestingly, the change response may gradually recover as the coma patient regains consciousness (Supplementary Fig. 6), indicating that the transitional click train paradigm could potentially monitor the entire recovery process of coma patients. However, no correlation was found between the CRS-R score, a standard method for quantifying the degree of coma, and either onset or change responses (Supplementary Fig. 7).

## Discussion

Our study meticulously examined the mechanisms of temporal merging within auditory perception, elucidating how the human auditory system assimilates discrete sound elements into unified auditory objects. With temporal merging, a click-train with minimal inter-click intervals (ICIs) gives a distinct auditory experience. Specifically, regular click trains (Reg_4-4.06_) prompted more pronounced change responses in the auditory cortex than irregular click trains (Irreg_4-4.06_), highlighting the significant impact of temporal regularity on auditory processing (Fig. 1). Further analysis demonstrated that the change response is intricately tied to the integration of multiple intervals, suggesting it as a marker for the perceptual transition between distinct auditory objects via temporal merging (Fig. 2). This response is notably affected by several factors: the length of ICI (Figs. 3a-c), the ICI ratio (IC1_2_ vs. ICI_1_) (Figs. 3d-f), and the regularity of the click train (Figs. 3g-i). Additionally, behavioral experiments showed enhanced change detection rates for regular click trains (Reg_4-4.06_) compared to irregular click trains and pure tones, corroborated by stronger EEG change responses (Fig. 4). Temporal continuity significantly affected behavioral and EEG responses, with better performance and clear tuning curves for continuous click trains compared to those with gaps (Fig. 5). Finally, the GFP method effectively distinguished coma patients from healthy subjects, suggesting the potential clinical application of transitional click trains for diagnosing and monitoring recovery in impaired consciousness (Fig. 6).

### Change response in transitional click train as a marker of temporal integration

Click trains with inter-click intervals (ICIs) less than approximately 33 ms are often perceived as pitch [21], and it has been suggested that the analysis of regularity in click trains differs for ICIs above and below 40–60 ms [22]. However, the neural correlates of pitch perception elicited by click trains remain under investigation. Auditory research utilizing click trains as stimuli has unveiled intricate neuronal responses in the auditory system on both single-neuron and systems neuroscience level. On single-neuron level, neurons display a remarkable capability for precise temporal coding where individual spike activities precisely align with specific intervals between the clicks [23]. Despite the prominence of this temporal alignment, rate coding emerges as another vital mechanism, particularly at accelerated click rates [24]. Lu et al.[7] identified two distinct populations of neurons: one that synchronizes to slow sound sequences and another that encodes rapid events through firing rates. However, these studies mainly focused on how individual clicks within a train are represented, largely overlooking the holistic perception of the click train as a coherent object [25]. At the macroscopic level, click trains have been extensively used to study auditory steady-state responses (ASSR) [26–28], where the neural response follows the same frequency of auditory stimuli, and the auditory response can be disrupted by an additional click [9]. These studies, similar to those at the single-neuron level, concentrate on responses to individual clicks, leaving the mechanism of how the brain integrates regular clicks into pitch perception unresolved. Recently, the holistic representation of sound has been investigated in the frequency domain, and researchers have found that auditory cortex (AC) neurons may exhibit bursting responses specifically to the configuration of tones but not to any constituent tone [29, 30]. However, the holistic representation in the temporal domain, especially for sound through temporal integration, has been seldom addressed. This gap exists because disentangling neural responses to individual clicks from those induced by the holistic perception of the whole click train as pitch poses a significant challenge. Consequently, no brain signal has yet been identified that adequately represents auditory events through temporal merging with click trains in prior research [7, 23, 24], highlighting a crucial area for future investigation at both single-neuron and macroscopic levels.

To navigate this intricacy, we propose the innovative concept of a transitional train, as illustrated in Figure 1. A typical onset response to transitional trains was observed in the first 300 ms of EEG signals, followed by an adaptation period from approximately 300 to 1000 ms, during which no discernible auditory response to individual clicks or the train was detected (Supplementary Fig. 2a). However, the introduction of a second click train with a slightly changed inter-click interval (ICI) (e.g., Reg_4-4.06_) within a transitional train elicited a change response in the adapted auditory cortex, followed by subsequent adaptation (Fig. 1d). Moreover, the transitional train also introduced a perceptual switch psychologically (Fig. 4). Since this change response in the EEG signal is not solely attributed to local temporal changes but is linked to temporal merging (Fig. 2), it most likely reflects a perceptual switch, signifying a transition between distinct temporal-merging auditory objects (Fig. 4). The key aspect underlying the transitional click train is that it maintains the presentation of individual clicks, which leads to consistent adaptation of the auditory cortex (Fig. 1), while simultaneously introducing a perceptual switch (Fig. 4). Therefore, the transitional train offers a convenient approach to investigate temporal integration. This innovative method allows us to disentangle the neural representation of individual clicks from that of the holistic auditory event, shedding light on the intricate process of temporal merging in auditory perception.

Three key factors influence the change response: the length of the inter-click interval (ICI) (Figs. 3a–c), the ICI ratio (ICI₂ vs. ICI₁) (Figs. 3d–f), and the regularity of the click train (Figs. 3g–i). Our research found that when the ICI length exceeds 32 ms, the auditory system is unable to elicit the change response, potentially suggesting a limit of temporal merging in ICI. This threshold is notably lower than what psychological studies have suggested, where the perception of a unified sound begins to falter at ICIs greater than 29.6 ms (Supplementary Fig. 1), and it is also below the 33 ms threshold often associated with pitch perception [21]. This discrepancy might be attributed to the inherent limitations of EEG recordings, which typically have a poor signal-to-noise ratio, underscoring the need for further investigation into the ICI threshold for temporal merging using more sophisticated methodologies. The auditory brain exhibits hypersensitivity to ICI ratios; even a 0.5% difference (Reg_4–4.02_) can evoke a robust change response (Figs. 3d–f). Meanwhile, the superior resolution of ICIs relative to pure tones (Fig. 4) suggests the involvement of a dedicated neuronal circuit with fine temporal resolution in temporal merging. Additionally, the regularity of the click train, which not only characterizes the temporal structure but also requires extended time for integration to extract the train’s regularity, reflects context-dependent temporal merging (Figs. 3g–i).The transitional click train paradigm presents significant opportunities for fundamental research in auditory science. Traditional auditory research has predominantly concentrated on the frequency domain, guided by the auditory system’s tonotopic organization, where distinct frequencies are processed separately along the auditory pathway [1]. Nevertheless, the importance of the temporal dimension in auditory processing cannot not be stressed enough. This temporal aspect is critical for speech and music perception, as well as for distinguishing environmental sounds [3]. Recent advances in neuroimaging and electrophysiology have enhanced our understanding of temporal integration mechanisms in oral language, revealing a hierarchical structure of temporal integration in the human brain [31, 32]. However, there remains a significant gap in our understanding of temporal integration in non-human animals, primarily due to the lack of a clear neuronal signature for this process, which has impeded research at the neuronal level and in animal studies. The identification of a change response in transitional click trains in our study provides a promising pathway to investigate this complex area further. Future research could employ the transitional click train paradigm to delve into the neuronal mechanisms underpinning temporal integration at the neuronal level in animal subjects.

In addition to providing signals for temporal integration, our study elucidates the neuronal mechanisms underlying pitch perception evoked by click trains. Our findings highlight the role of temporal integration as a key process in pitch perception. Traditional theories distinguish between resolved and unresolved harmonics based on the auditory system’s ability to segregate individual harmonic components [33, 34]. Resolved harmonics arise from distinct components processed by separate auditory filters, while unresolved harmonics involve closely spaced components processed by a single filter, relying on temporal coding for pitch extraction. Interestingly, sounds with the same repetition rate but very different spectral compositions often evoke the same pitch, while sounds with similar spectra can produce significantly different pitches. This observation demonstrates that the frequency-to-place mapping performed by the cochlea does not necessarily correspond to a frequency-to-pitch mapping [35]. Temporal pitch, induced by click trains, is distinct in that it relies solely on the temporal regularity of successive auditory events rather than on the spectral components [35–38]. Our study provides compelling neuronal evidence supporting this process, demonstrating that the change response reflects the integration of temporal information into a unified auditory pitch (Fig. 2). Previous research has used transitional click trains to investigate temporal pitch sensitivity [39] and observed the change response in EEG signals of cats [38]. Our insertion experiments further explored the nature of the change response, with a focus on temporal integration (Fig. 2).

### Behavior relevance during transitional click train

The alignment between psychological findings and EEG data underscores a notable facet of our research. On the one hand, our psychological data reveal the heightened sensitivity of regular transitional click trains compared to both pure tones (Figs. 4e, g, i) and irregular click trains (Figs. 4d, f, h). Concurrently, EEG signals exhibit stronger responses to regular click trains (Figs. 4f–i). Regular click trains, especially those with shorter inter-click intervals (ICIs), are often perceived to have pitch-like qualities. This perception has traditionally been explained through theories and computational models focusing on the basilar membrane’s processing [10]. The pronounced sensitivity of regular click trains over pure tones underscores the critical role of temporal integration in the central nervous system for refining fine temporal structures. This suggests that the pitch perception associated with regular click trains might originate in the central auditory system rather than the basilar membrane. This hypothesis necessitates further exploration, particularly employing our innovative transitional click train in animal studies.

Additionally, both psychological and EEG responses demonstrate a dependency on temporal continuity. The introduction of a 600-ms gap adversely affects change detection capabilities (Figs. 5b, c) and alters the tuning of change differences (Figs. 5f, g). Given that the change response to the transitional click train systematically correlates with the contrast ratio (Figs. 5f, g), whereas responses to the second sound in the gap condition remain relatively constant across different contrast ratios, it suggests that the change response primarily signifies the signal of perceptual switching rather than the perception of the second pitch. The influence of temporal discontinuity on both behavioral and neural responses accentuates the essential role of temporal integration within the auditory system, suggesting that seamless auditory perception relies on the continuous flow of temporal information.

### Change response as a biomarker in clinical application

Three key factors influence the change response (Fig. 3) and are consequently related to temporal merging, offering diverse metrics for characterizing temporal integration, and potentially serving as valuable tools in clinical applications: the length of the ICI, the difference between ICIs, and the regularity in the click train. These factors hold promise as potential biomarkers for mental disorders. To further explore this possibility, we investigated the coma patients (Fig. 6). The change response dramatically vanished (Fig. 6a), even in some cases, the onset response exists while no visible change response (Supplementary Fig. 5c), suggesting of different origin of both kinds of response. Interestingly, the change response may recover as the patient get recovery (Supplementary Fig. 6). The observed change response to transitional train provides an innovative pathway for refining coma monitoring techniques. Further extending the clinical applicability of our research, we propose the use of transitional trains for the assessment of psychiatric conditions. As temporal integration, a central component of brain functionality [12, 13], has been found to be compromised in conditions like schizophrenia [14], autism spectrum disorders [15, 16], attention deficit hyperactivity disorder [17], and Parkinson’s Disease [18]. Given these findings, the signal of temporal integration might be poised to emerge as a pivotal biomarker for broader clinical diagnostics.

## Methods

### Experimental Procedure and Participants

The study comprised four experiments, all conducted in accordance with the Declaration of Helsinki (2013) [40]. Experiments 1-3 were conducted with healthy participants (48 males and 40 females in total, mean age: 25.17 years, standard deviation: 5.15). Participants maintained a stationary head position while listening to auditory stimuli and responding via keyboard presses. These experiments were approved by the Institutional Review Board (IRB-20230131-R), and informed consent was obtained from all participants. Experiment 4 involved 22 coma participants with impaired consciousness (16 males and 6 females, mean age: 56.52 years, standard deviation: 15.96), including one participant who was recorded again after recovery from coma. The level of consciousness in coma participants was measured using the Coma Recovery Scale-Revised (CRS-R) scores. This experiment was approved by Natural Science Foundation of Zhejiang Provincial (LGF22H170006).

**Experiment 1**: This was a gap detection task (Supplementary Fig. 1a) involving click trains of 1024 ms duration with varying ICIs (4, 8, 16, 32, 64, 128, 256 ms). Participants were positioned in a chair facing a keyboard and speaker. After each click train, a 100-ms 1000-Hz cue (100 ms) was presented 800 ms after the end of the click train. Participants were instructed to press the right key on the keyboard if a gap was detected in the click train and the left key if the click train was perceived as continuous. Keyboard press was valid within 700 ms after the cue onset.

**Experiment 2**: This included two passive listening sessions followed by two discrimination sessions.

- **Passive Listening Session 1**: Four regular transitional trains (Fig. 3a) were randomly presented (Reg_4-4.06_, Reg_8-8.12_, Reg_16-16.24_, Reg_32-32.48_).
- **Passive Listening Session 2**: The session (Fig. 3d) consisted of five regular transitional trains (Reg_4-4_, Reg_4-4.01_, Reg_4-4.02_, Reg_4-4.03_, Reg_4-4.06_) and two irregular transitional trains (Irreg_4-4_, Irreg_4-4.06_), and a tone-pair (Tone_250-246_).
- **Discrimination Session 1**: The session (Fig. 4a) included stimuli from Passive Listening Session 2, another irregular transitional train (Irreg_4-8_), and another tone-pair (Tone_250-250_). Participants needed to report whether the sound changed by pressing one of two designated keys, with left key representing change in the transitional stimulation and right key for no change.
- **Discrimination Session 2**: A 600-ms gap between click-train 1 and click-train 2 resulted in gap transitional trains (Fig. 5a). Participants identified if the two click-trains were changed and report following the same rule in Discrimination Session 1.

**Experiment 3** consisted of two passive listening sessions.

- **Passive Listening Session 1**: 1, 2, 4, 8, 16, and 32 intervals (each with an interval of 4.06 ms) were inserted into a click train with a 4 ms ICI (Fig. 2a). Controls included Reg_4-4_ and Reg_4-4.06_.
- **Passive Listening Session 2**: Four Irreg_4-4.06_ transition trains with different standard deviations and Reg_4-4.06_ were randomly presented (Fig. 3g).
- **Passive Listening Session 3**: Two transitional trains, Reg_4-4_ and Reg_4-5_, were randomly presented. This session was added as a control for the coma patients.

**Experiment 4** involved one passive listening session with Reg_4-4_ and Reg_4-5_ transitional trains presented to coma participants.

## Auditory Stimuli

The experiments were conducted within a sound-proof room. A single click consisted of a 0.2-ms pulse. Click trains were categorized as either regular, with a fixed ICI, or irregular, with random ICIs. For irregular click trains, the ICIs were randomized using a Gaussian distribution, and satisfied the following formula:

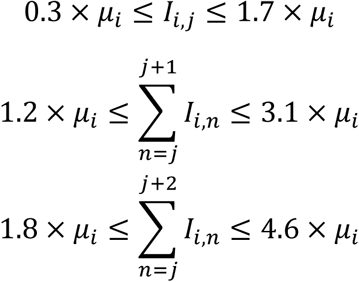

here, μ_*i*_ represents the average ICI of the *i*^th^ train in the transitional click train and *I*_*i*,*j*_ is the *j*^th^ ICI in the *i*^th^ train. The mean value of the Gaussian distribution matched the fixed ICI of a regular click train, while the standard deviation was a certain percentage of the fixed ICI (0.25%, 0.5%, 1%, or 50%). A transitional click train is formed by concatenating two click trains. For example, a Reg_4-4.06_ transitional train denotes the combination of two regular click trains: regular click train 1 (with an ICI of 4 ms) is seamlessly followed by regular click train 2 (with an ICI of 4.06 ms). Similarly, an irregular transitional train is composed of two irregular click trains with the given average ICIs. For continuous click trains that seamlessly transitioned from ICI_1_ to ICI_2_, the transition time was defined as the onset time of the first click after the first ICI_2_ interval (Fig. 1b). The ratio of ICI_1_ to ICI_2_ quantified the difference level between the two ICIs. Auditory stimuli were delivered through the Golden Field M23 sound player, driven by a Creative AE-7 Sound Blaster, with a sampling rate of 384 kHz. Sound delivery was controlled with Psychtoolbox 3 in MATLAB. Sound intensity was calibrated to maintain a constant level of 60 dB SPL (sound pressure level), using a ¼-inch condenser microphone (Brüel & Kjær 4954, Nærum, Denmark) and a PHOTON/RT analyzer (Brüel & Kjær, Nærum, Denmark).

## Data Acquisition

In Experiment 2, EEG data were acquired using a 64-channel NeuroScan system (Compumedics, USA). Experiments 3 and 4 utilized a 64-channel NeuSenW system (Neuracle, China). The EEG data were sampled at 1 kHz, and electrode placement followed the international 10-20 system protocol.

## Data Analysis

The data analyses were performed using MATLAB R2021b (MathWorks) and the Fieldtrip toolbox. Monopolar referencing was employed in this study. The multichannel EEG data underwent several preprocessing steps. First, the full EEG data were filtered using a band-pass filter in the frequency range of 0.5 to 40 Hz. Then, 4-second epochs were obtained, spanning from –1 to 3 seconds relative to trial onset. Independent component analysis (ICA) was then applied to the epochs to remove electrooculogram (EOG). After ICA, baseline correction was applied by subtracting the mean response within the baseline window from –200 to 0 ms relative to the onset of train stimulation for each trial. Following this, a relative threshold was used to evaluate motion artifacts for each trial, excluding those exceeding predefined thresholds. The relative threshold was determined based on the percentage of bad samples within a trial. A sample was flagged as a “bad sample” if it fell outside the range of “Mean”±3×“SD” for a trial across specific channels. Trials with over 20% of bad samples were labeled as “bad trials”. Additionally, channels with over 10% of bad trials were labeled as “bad channels”. Bad channels were initially excluded by assessing bad samples using data from all channels, and subsequently, bad trials were excluded from all channels by computing bad samples using data from the remaining good channels. Finally, the event-related potential (ERP) data were obtained by averaging the epoch data for each experimental condition, channel, and subject. Prior to applying inter-subject analysis, ERP data were normalized by each channel’s standard deviation per subject to reduce inter-channel variability.

For time-sample level comparisons of ERP or global field power (GFP) between conditions, a two-tailed cluster-based permutation test was conducted using the ‘ft_timelockstatistics’ function from the FieldTrip toolbox in MATLAB.

The change response comprised two major ERP components, cP1 and cN2 (Supplementary Fig. 2c and Fig. 1d), which exhibited opposite polarities in the frontal and temporal-parietal-occipital scalp regions (Supplementary Fig. 2a). To quantify individual components (e.g., cP1 of Reg_4-4.01_ in Supplementary Fig. 3, and P1/cP1/N2/cN2 in Figs. 6f, g), we calculated the mean ERP value within a [-25, 25] ms time window centered around the peak time, subtracting the baseline mean ([-200, 0] ms relative to change or onset). For a global quantification of the entire change or onset response across the scalp, we used the relative response magnitude (RM), defined as the root mean square (RMS) of the ERP over a specific time window encompassing both peak and trough responses, with the baseline RMS subtracted. The RM calculation proceeded as follows:

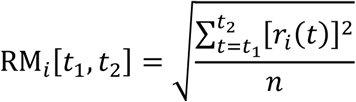

here, r_*i*_(*t*) represents the ERP of the *i*^*t*ℎ^ channel at time *t*, *n* represents the number of samples within the time window from *t*_1_ and *t*_2_, and RM_*i*_ is the response magnitude of channel *i*. The relative RM was defined as:

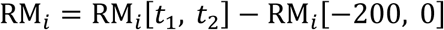

The time window for the change response was determined using a two-tailed cluster-based permutation test of the GFP data between the transitional click trains Reg_4-4.06_ and Reg_4-4_ at the time-sample level (Supplementary Fig. 2c). Specifically, the change response time windows were [74, 251] ms relative to change for all passive listening sessions. Channels with significant change responses were tested with a two-tailed paired t-test (for N>30) or a Wilcoxon signed rank test (for N<30) between the RM of change and the RM of baseline at subject level for each channel. The relative RMs of all channels were then averaged for each subject to facilitate comparison across experimental conditions (e.g., tunings and scatterplots).

To quantify change responses in subjects with impaired consciousness, global field power (GFP) was employed due to its robustness to spatial variability and enhanced sensitivity to response latency. GFP calculates the differences in potential across all electrodes at each sampling point, thereby mitigating the influence of spatial variability in electrode placement or individual differences in brain anatomy. The calculation of GFP follows:

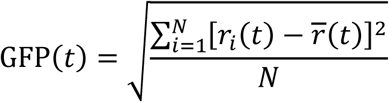

here, *N* is the total number of channels, r_*i*_(*t*) represents the ERP of the *i*^*t*ℎ^ channel at time *t*, and r(*t*) denotes the averaged response across all channels at time *t*. The difference between the maximum GFP value detected within 300 ms after the change point of the train stimulation and the mean GFP value across the 200-ms baseline response before change was used as an indicator of the change response.

The ratio of change detection for each group was calculated by dividing the number of trials in which the subject pressed the left arrow key (change detection) by the total number of trials in that group. In total, 36 subjects were included in the stand-alone analyses of no-gap change detection task. In the comparison of gap and no-gap change detection tasks, only the intersection of subjects, which consisted of 34 subjects, was included. Psychometric functions were fitted to data using a cumulative Gaussian function [41, 42]:

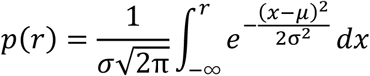

here, *p*(r) represents the ratio of change detection as a function of ICI r, μ is the Gaussian mean, and σ is the standard deviation (SD). The threshold of change detection was defined as 0.6 of the Gaussian fit (Fig. 5c). This curve fitting procedure was achieved using ‘psignifit’ software package (see http://bootstrap-software.org/psignifit/) for MATLAB. Similarly, the ICI threshold of the ratio of gap detection (Supplementary Fig. 1b) was obtained with the same fitting procedure.

## Figure Legend

**Supplementary Figure 1.**
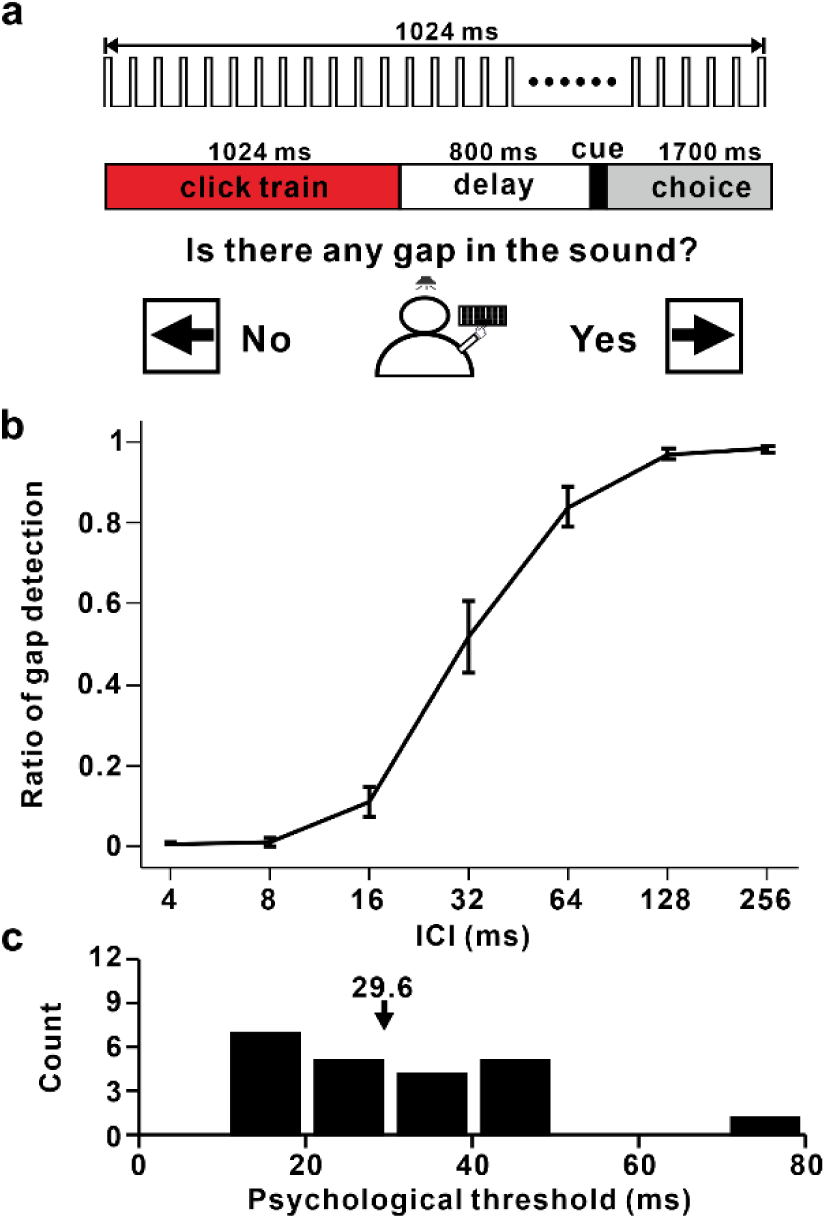
Gap detection task in click train sequences. (**a**) Schematic of the gap detection task. Participants were seated in a quiet room, facing a computer screen with a keyboard. Click trains were played through a speaker, each lasting 1024 ms with inter-click intervals (ICIs) varying from 4 ms to 256 ms. Following each click train, a cue tone appeared after 800 ms, prompting participants to press the right key within 1700 ms if they detected a gap, or the left key if the click train was perceived as a continuous sound. (**b**) Group Psychometric Function: This plot aggregates the responses from all participants (n=22). The horizontal axis shows the ICIs of the click trains, while the vertical axis represents the frequency of right key presses, indicative of gapped sound perception. Error bars denote the standard error (SE) of the mean. (**c**) Distribution of the psychological thresholds for gap detection among the participants: the arrow indicates the mean threshold value across the participants.

**Supplementary Figure 2.**
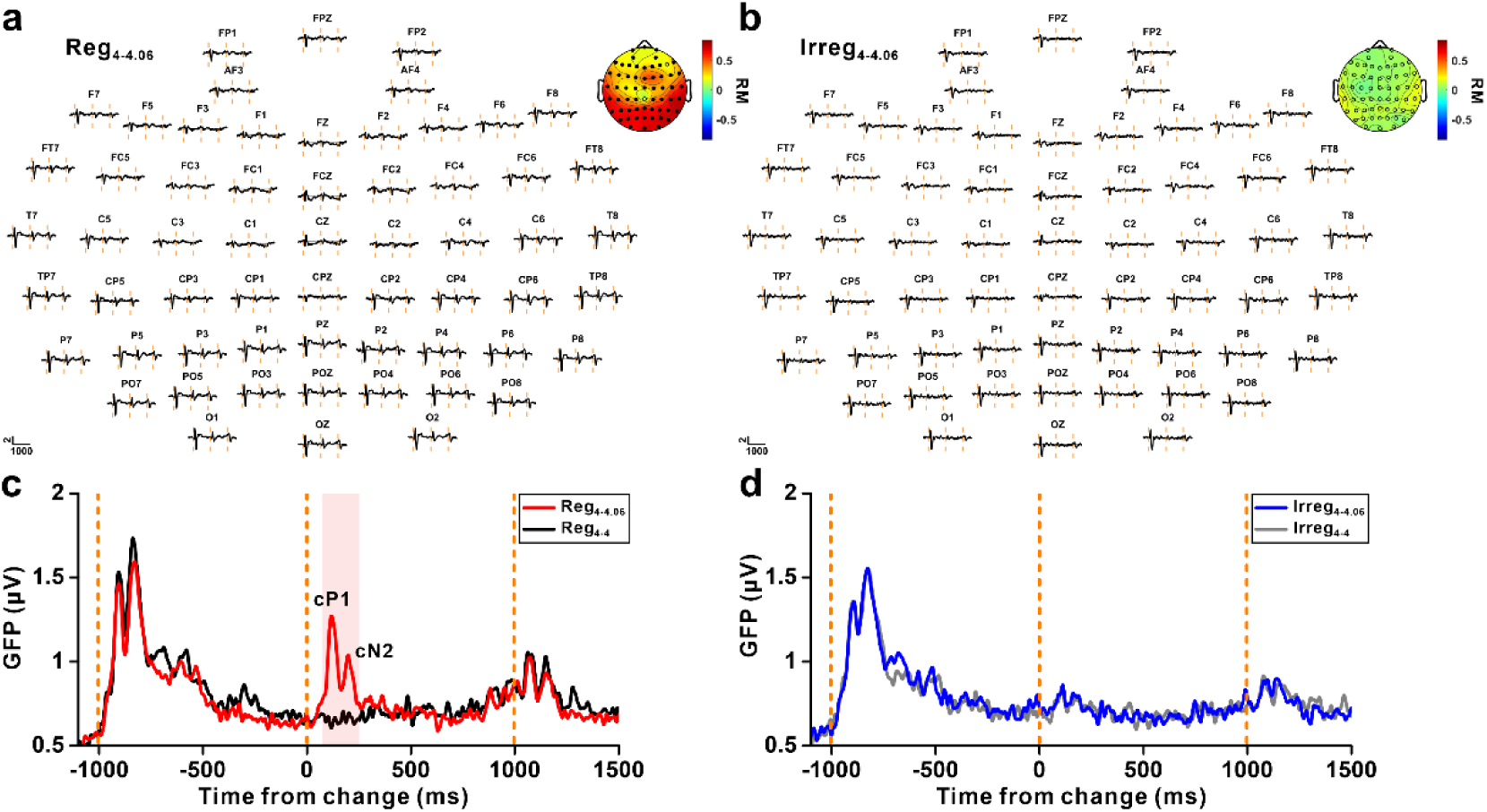
Change responses in transitional click trains in 64-channel EEG. (**a-b**) Multichannel event-related potentials (ERPs) for transitional click trains Reg_4-4.06_ (a) and Irreg_4-4.06_ (b), with orange dashed lines indicating onset, change, and offset from left to right, respectively. Topographic maps (top right) display the relative response magnitude (RM) for each channel. Channels with significant change responses are marked with filled circles (p<0.05, paired t-test). (**c-d**) Global field power comparisons: Reg_4-4.06_ vs. Reg_4-4_ (c) and Irreg_4-4.06_ vs. Irreg_4-4_ (d). The red bar in (c) indicates the time window [74, 251] ms, highlighting significant GFP difference between Reg_4-4.06_ and Reg_4-4_ (two-tailed permutation test).

**Supplementary Figure 3.**
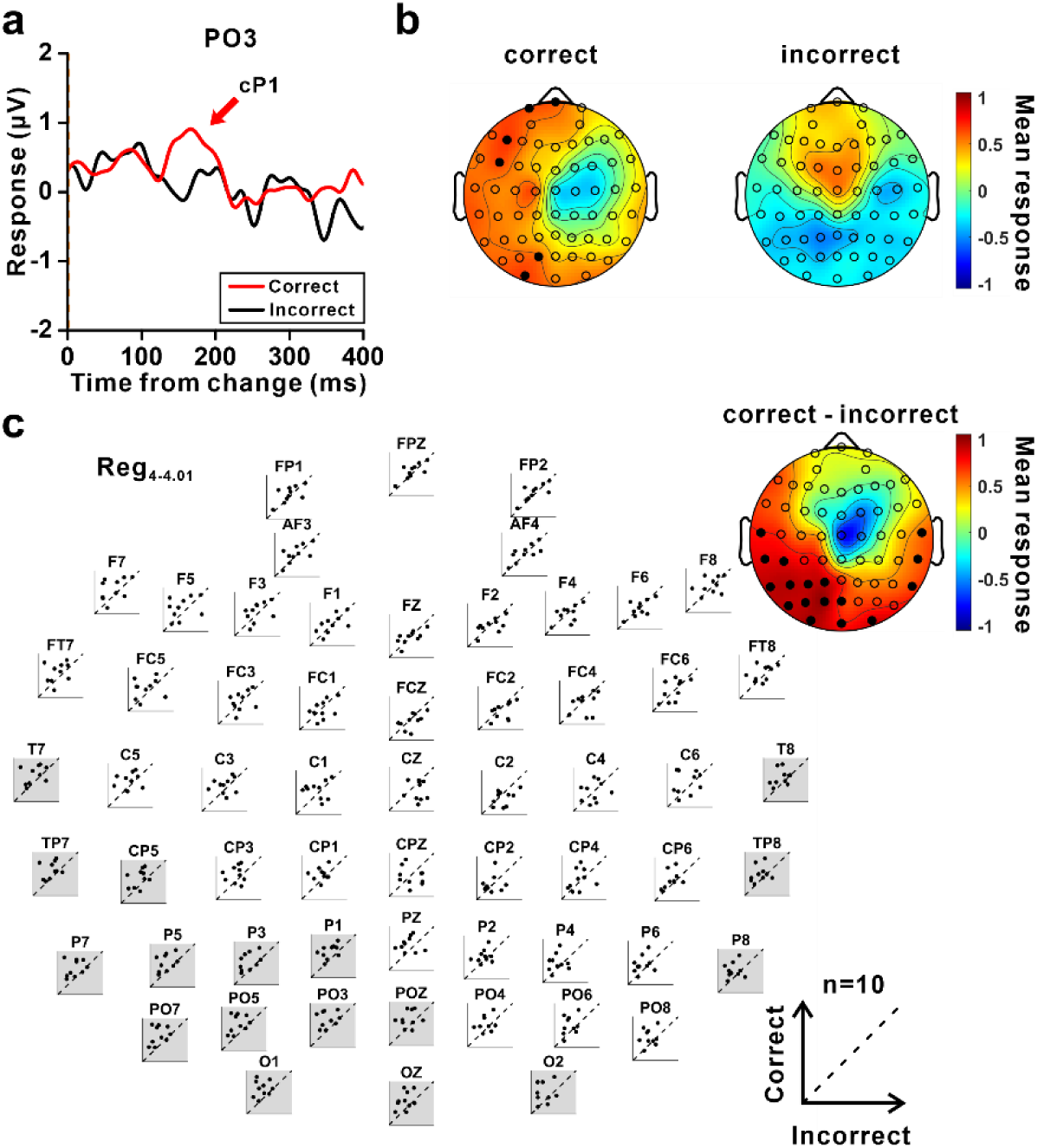
cP1 component correlates with cognitive processing. (**a**) Average waves at PO3 for correct (red) and incorrect (black) trials under Reg_4-4.01_. Subjects included in this analysis had a change detection ratio between 0.3 and 0.7 for Reg_4-4.01_ (N=10). (**b**) Topographic maps of average cP1 responses (peaked at 163 ms relative to change) for correct (left) and incorrect (right) trials under Reg_4-4.01_, with filled circles indicating channels with cP1 responses significantly different from baseline (p<0.05, Wilcoxon signed rank test). (**c**) Scatterplots of average cP1 responses between correct and incorrect trials for each channel under Reg_4-4.01_, with a differential topographic map in the top right. Channels with significant differences between correct and incorrect trials are highlighted with gray backgrounds in scatterplots and filled circles in the map (p<0.05, Wilcoxon signed rank test).

**Supplementary Figure 4.**
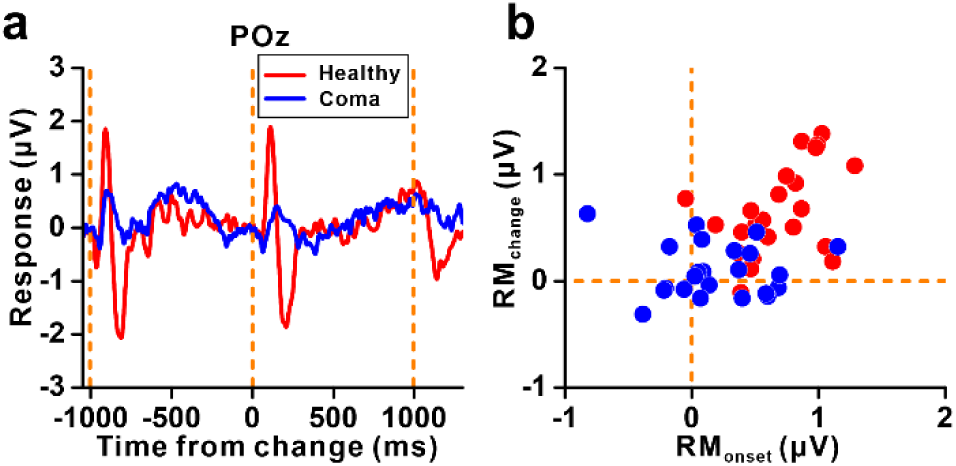
The effect of consciousness on change responses in transitional train, indexed with ERP. (**a**) Average waves at POz of coma participants (blue line, n=22) and healthy participants (red line, n=24) under Reg_4-5_. (**b**) Scatterplots of relative RM of change responses and onset responses of coma participants (blue) and healthy participants (red).

**Supplementary Figure 5.**
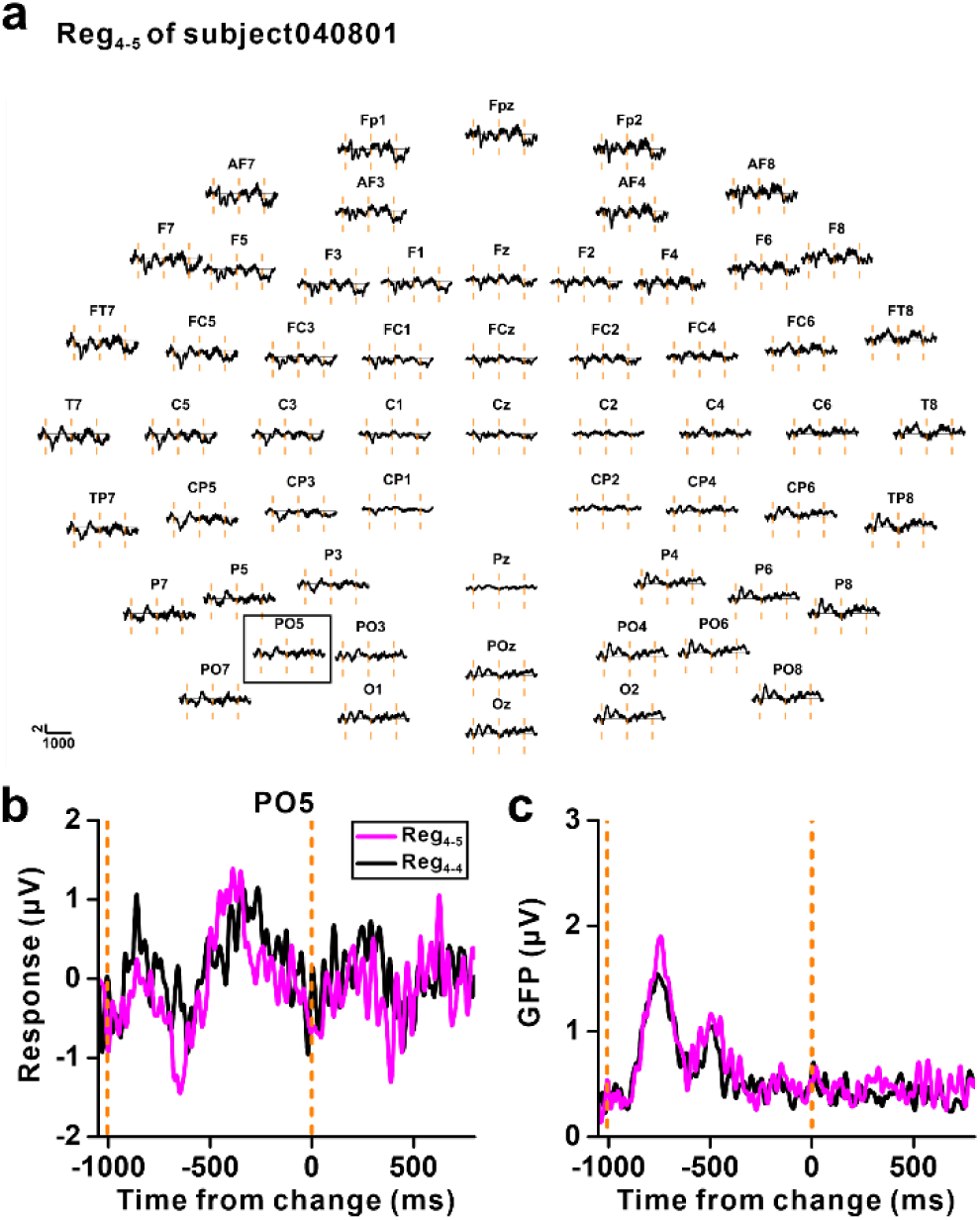
Spatial variability in EEG responses of coma participants. (**a**) Evoked onset and change responses under Reg_4-5_ in one coma participant. (**b**) Average waves at PO5 under Reg_4-5_ (magenta) and Reg_4-4_ (black). (**c**) Global field power under Reg_4-5_ (magenta) and Reg_4-4_ (black).

**Supplementary Figure 6.**
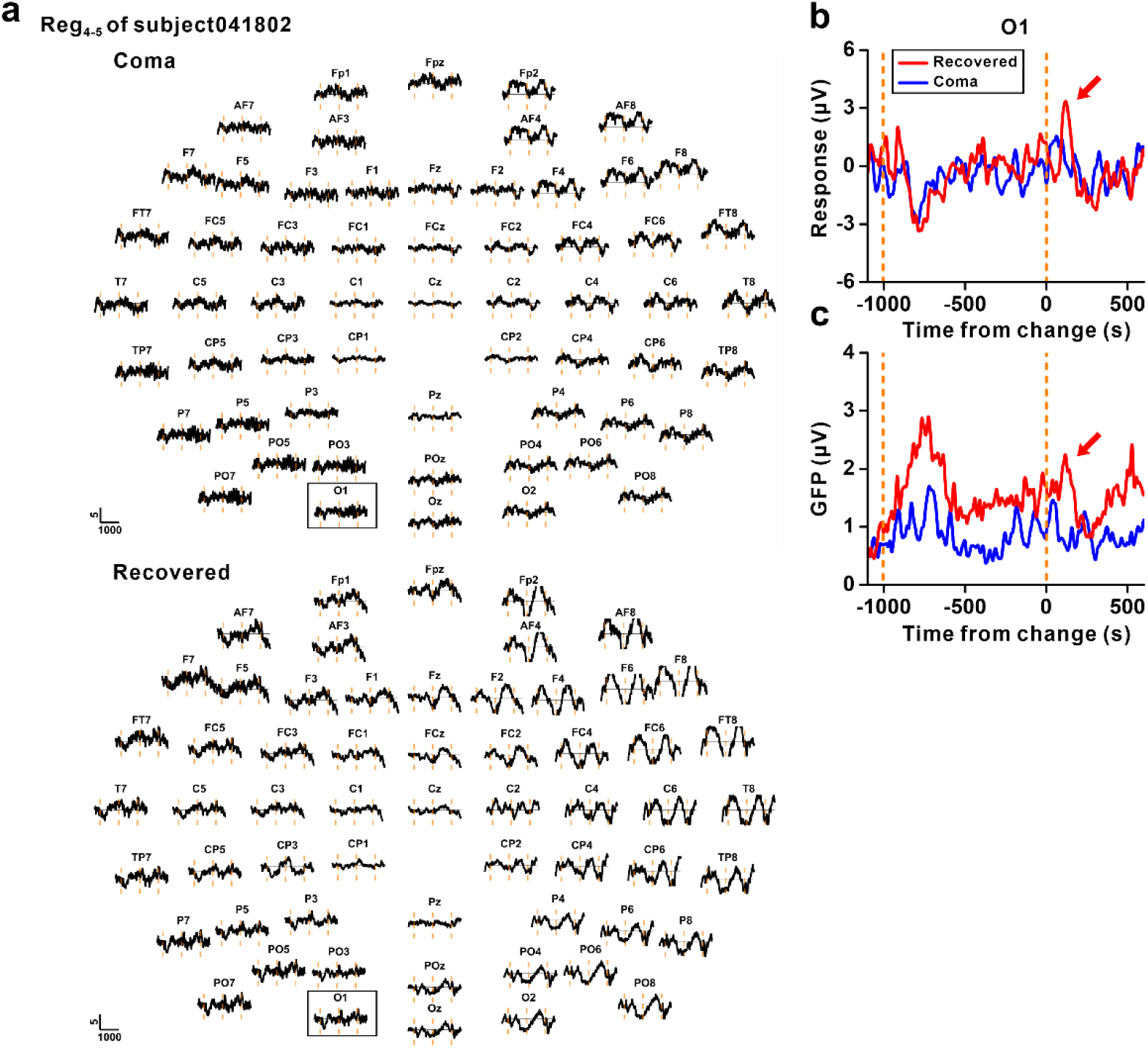
Regain of change responses after recovery from coma. (**a**) Evoked onset and change responses under Reg_4-5_ before (above) and after recovery (below) in one coma participant who recovered two months after the initial recording. The black blocks mark the O1 channel displayed in (b). (**b**) The evoked onset and change responses at O1 under coma (blue) and recovered (red) conditions. (**c**) Global field power under coma (blue) and recovered (red) conditions.

**Supplementary Figure 7.**
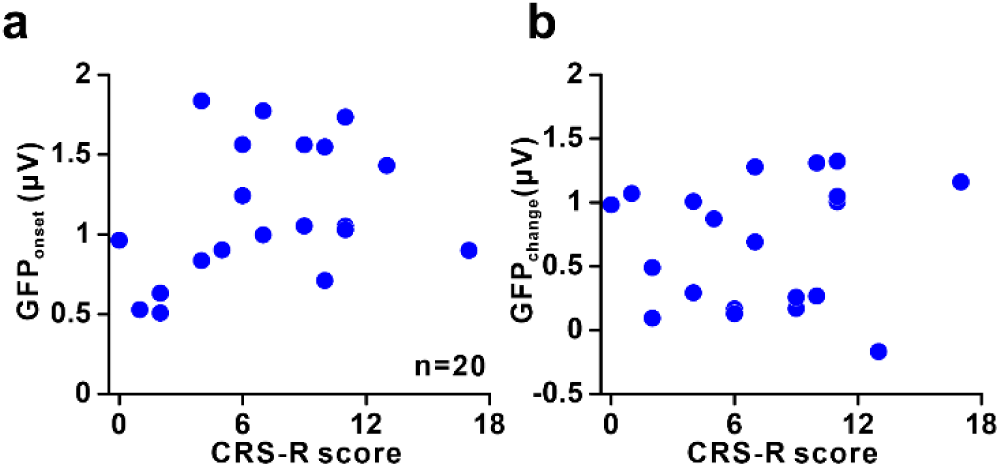
Onset and change responses are not consistent with CRS-R scores. (**a**) Scatterplots showing GFP indices of onset responses plotted against CRS-R score (n=20). (**b**) Scatterplots showing GFP indices of change responses plotted against CRS-R score. Both show no significant correlation with CRS-R scores: Pearson’s correlation, p=0.18 for onset responses and p=0.66 for change responses.

## Author contributions

Conceptualization, X.Y.; Methodology, X.Y., P.S., H.X. and H.Y.; Investigation and Data Curation, X.Y., P.S., H.X., H.Y., X.D., Y.Z., X.B., Q.H., I.M., H.T., W.N., Z.T., P.C., T.Z., L.Z. and X.Z.; Writing – Original Draft, X.Y., P.S., H.X. and H.Y.; Writing – Review & Editing, X.Y., X.Z., P.S., H.X., H.Y., and L.C.; Funding Acquisition, X.Y., and Y.Z.; Resources, X.Y.; Software, P.S., H.X. and H.Y.; Validation and Visualization, P.S., H.X. and H.Y.; Supervision, X.Y.; L.C.; W.W. and X.Z.

## Acknowledgments

We are grateful to Xiaokai Kou and Fujin Gao and for their help with the experiments. This work was supported by STI2030-Major Projects (2022ZD0204800 and 2022ZD0204600) (to X.Y.); National Natural Science Foundation of China 32171044 (to X.Y.), 32100827 (to Y.Z.), and 32271078 (to L.C.); Key Support Discipline Construction Project of Shanghai Municipal Health Commission 2023ZDFC0203 (to X.Z.); Zhejiang Provincial Natural Science Foundation of China under Grant No. LGF20H020010 (to W.W.); Natural Science Foundation of Zhejiang Provincial LGF22H170006 (to L.Z.)

## References

1. Schnupp, J., I. Nelken, and A.J. King, Auditory Neuroscience: Making Sense of Sound. 2010: Auditory Neuroscience: Making Sense of Sound.

2. ten Cate, C. and M. Spierings, Rules, rhythm and grouping: auditory pattern perception by birds. Animal Behaviour, 2019. 151: p. 249–257.

3. Moore, B.C.J., Temporal integration and context effects in hearing. Journal of Phonetics, 2003. 31(3-4): p. 563–574.

4. Gao, R., et al., Neuronal timescales are functionally dynamic and shaped by cortical microarchitecture. Elife, 2020. 9.

5. Ding, N., et al., Cortical tracking of hierarchical linguistic structures in connected speech. Nat Neurosci, 2016. 19(1): p. 158–64.

6. Norman-Haignere, S.V., et al., Multiscale temporal integration organizes hierarchical computation in human auditory cortex. Nat Hum Behav, 2022. 6(3): p. 455–469.

7. Lu, T., L. Liang, and X. Wang, Temporal and rate representations of time-varying signals in the auditory cortex of awake primates. Nat Neurosci, 2001. 4(11): p. 1131–8.

8. Steinschneider, M., et al., Click train encoding in primary auditory cortex of the awake monkey: evidence for two mechanisms subserving pitch perception. J Acoust Soc Am, 1998. 104(5): p. 2935–55.

9. Lutkenhoner, B. and R.D. Patterson, Disruption of the auditory response to a regular click train by a single, extra click. Exp Brain Res, 2015. 233(6): p. 1875–92.

10. Balaguer-Ballester, E., S.L. Denham, and R. Meddis, A cascade autocorrelation model of pitch perception. J Acoust Soc Am, 2008. 124(4): p. 2186–95.

11. Yost, W.A., et al., Pitch strength of regular-interval click trains with different length “runs” of regular intervals. J Acoust Soc Am, 2005. 117(5): p. 3054–68.

12. Mauk, M.D. and D.V. Buonomano, The neural basis of temporal processing. Annu Rev Neurosci, 2004. 27: p. 307–40.

13. D’Argembeau, A., et al., The neural basis of temporal order processing in past and future thought. J Cogn Neurosci, 2015. 27(1): p. 185–97.

14. Su, L., et al., Temporal perception deficits in schizophrenia: integration is the problem, not deployment of attentions. Sci Rep, 2015. 5: p. 9745.

15. Stevenson, R.A., et al., Multisensory temporal integration in autism spectrum disorders. J Neurosci, 2014. 34(3): p. 691–7.

16. Nakano, T., et al., Deficit in visual temporal integration in autism spectrum disorders. Proc Biol Sci, 2010. 277(1684): p. 1027–30.

17. Panagiotidi, M., P.G. Overton, and T. Stafford, Multisensory integration and ADHD-like traits: Evidence for an abnormal temporal integration window in ADHD. Acta Psychol (Amst), 2017. 181: p. 10–17.

18. Tokushige, S.I., et al., Does the Clock Tick Slower or Faster in Parkinson’s Disease? – Insights Gained From the Synchronized Tapping Task. Front Psychol, 2018. 9: p. 1178.

19. Skrandies, W., Global field power and topographic similarity. Brain topography, 1990. 3: p. 137–141.

20. Giannopoulos, A.E., et al., Early auditory-evoked potentials in body dysmorphic disorder: An ERP/sLORETA study. Psychiatry Research, 2021. 299: p. 113865.

21. Krumbholz, K., R.D. Patterson, and D. Pressnitzer, The lower limit of pitch as determined by rate discrimination. J Acoust Soc Am, 2000. 108(3 Pt 1): p. 1170–80.

22. Phillips, D.P., et al., Dual mechanisms in the perceptual processing of click train temporal regularity. J Acoust Soc Am, 2012. 132(1): p. EL22–8.

23. Oshurkova, E., H. Scheich, and M. Brosch, Click train encoding in primary and non-primary auditory cortex of anesthetized macaque monkeys. Neuroscience, 2008. 153(4): p. 1289–99.

24. Bendor, D. and X. Wang, Differential neural coding of acoustic flutter within primate auditory cortex. Nat Neurosci, 2007. 10(6): p. 763–71.

25. Song, P., et al., A new function of offset response in the primate auditory cortex: marker of temporal integration. Commun Biol, 2024. 7(1): p. 1350.

26. Nakamura, T., et al., Characteristics of auditory steady-state responses to different click frequencies in awake intact macaques. BMC Neurosci, 2022. 23(1): p. 57.

27. Presacco, A., et al., Auditory steady-state responses to 40-Hz click trains: relationship to middle latency, gamma band and beta band responses studied with deconvolution. Clin Neurophysiol, 2010. 121(9): p. 1540–1550.

28. Neklyudova, A., et al., Atypical brain responses to 40-Hz click trains in girls with Rett syndrome: Auditory steady-state response and sustained wave. Psychiatry Clin Neurosci, 2024.

29. Li, R., et al., Holistic bursting cells store long-term memory in auditory cortex. Nat Commun, 2023. 14(1): p. 8090.

30. Wang, M., et al., Single-neuron representation of learned complex sounds in the auditory cortex. Nat Commun, 2020. 11(1): p. 4361.

31. Chang, C.H.C., S.A. Nastase, and U. Hasson, Information flow across the cortical timescale hierarchy during narrative construction. Proc Natl Acad Sci U S A, 2022. 119(51): p. e2209307119.

32. Lerner, Y., et al., Topographic mapping of a hierarchy of temporal receptive windows using a narrated story. J Neurosci, 2011. 31(8): p. 2906–15.

33. Carlyon, R.P. and T.M. Shackleton, Comparing the fundamental frequencies of resolved and unresolved harmonics: Evidence for two pitch mechanisms? The Journal of the Acoustical Society of America, 1994. 95(6): p. 3541–3554.

34. Shackleton, T.M. and R.P. Carlyon, The role of resolved and unresolved harmonics in pitch perception and frequency modulation discrimination. The Journal of the Acoustical Society of America, 1994. 95(6): p. 3529–3540.

35. Plack, C.J., A.J. Oxenham, and R.R. Fay, Pitch: neural coding and perception. Vol. 24. 2006: Springer Science & Business Media.

36. Carlyon, R.P. and J.M. Deeks, Limitations on rate discrimination. The Journal of the Acoustical Society of America, 2002. 112(3): p. 1009–1025.

37. Macherey, O. and R.P. Carlyon, Re-examining the upper limit of temporal pitch. The Journal of the Acoustical Society of America, 2014. 136(6): p. 3186–3199.

38. Richardson, M.L., et al., Temporal pitch sensitivity in an animal model: Psychophysics and scalp recordings: Temporal pitch sensitivity in cat. Journal of the Association for Research in Otolaryngology, 2022. 23(4): p. 491–512.

39. Ungan, P. and S. Yagcioglu, Significant variations in Weber fraction for changes in inter-onset interval of a click train over the range of intervals between 5 and 300 ms. Frontiers in psychology, 2014. 5: p. 1453.

40. Duncan, N., Declaration of Helsinki. World Medical Journal, 2013. 59(4).

41. Yu, X.J., et al., Neuronal thresholds and choice-related activity of otolith afferent fibers during heading perception. Proc Natl Acad Sci U S A, 2015. 112(20): p. 6467–72.

42. Xu, X.X., et al., Adaptation facilitates spatial discrimination for deviant locations in the thalamic reticular nucleus of the rat. Neuroscience, 2017. 365: p. 1–11.

